# Understanding SARS-CoV-2 Spike glycoprotein clusters and their impact on immunity of the population from Rio Grande do Norte, Brazil

**DOI:** 10.1101/2023.10.05.561101

**Authors:** Diego Gomes Teixeira, João Firmino Rodrigues-Neto, Dayse Caroline Severiano da Cunha, Selma Maria Bezerra Jeronimo

**Affiliations:** Instituto de Medicina Tropical do Rio Grande do Norte, Universidade Federal do Rio Grande do Norte, Natal, Rio Grande do Norte, Brazil; Escola Multicampi de Ciências Médicas do Rio Grande do Norte, Universidade Federal do Rio Grande do Norte, Caicó, Rio Grande do Norte, Brazil; Biochemistry Department, Centro de Biociências, Universidade Federal do Rio Grande Norte, Natal, Rio Grande do Norte, Brazil; Instituto Nacional de Ciência e Tecnologia de Doenças Tropicais, Natal, Rio Grande do Norte, Brazil

**Keywords:** SARS-CoV-2 variants, Spike glycoprotein, epitope prediction, COVID-19, cytokine induction, immunoinformatics

## Abstract

SARS-CoV-2 genome underwent mutations since it started circulating intensively within the human populations. The aim of this study was to understand the fluctuation of the spike clusters concomitant to high rate of population immunity either due to natural infection and/or vaccination in a state of Brazil that had high rate of infection and vaccination coverage. A total of 1715 SARS-CoV-2 sequences from the state of Rio Grande do Norte, Brazil, were retrieved from GISAID and subjected to cluster analysis. Immunoinformatics were used to predict T- and B-cell epitopes, followed by simulation to estimate either pro- or anti-inflammatory responses and correlate with circulating variants. From March 2020 to June 2022, Rio Grande do Norte reported 579,931 COVID-19 cases with a 1.4% fatality rate across three major waves: May-Sept 2020, Feb-Aug 2021, and Jan-Mar 2022. Cluster 0 variants (wild type strain, Zeta) were prevalent in the first wave and Delta in the latter half of 2021, featuring fewer unique epitopes. Cluster 1 (Gamma [P1]) dominated the first half of 2021. Late 2021 had Clusters 2 (Omicron) and 3 (Omicron sublineages) with the most unique epitopes, while Cluster 4 (Delta sublineages) emerged in the second half of 2021 with fewer unique epitopes. Cluster 1 epitopes showed a high pro-inflammatory propensity, while others exhibited a balanced cytokine induction. The clustering method effectively identified Spike groups that may contribute to immune evasion and clinical presentation, and explain in part the clinical outcome.

**IMPORTANCE:** Identification of epitopes of emerging or endemic pathogens is of importance to estimate population responses and predict clinical outcomes and contribute to vaccine improvement. In the case of SARS-CoV-2, the virus within 6 months of circulation transitioned from the wild-type to novel variants leading to distinct clinical outcomes. Immunoinformatics analysis of viral epitopes of isolates from the Brazilian state of Rio Grande do Norte was performed using a clustering method. This analysis aimed to clarify how the introduction of novel variants in a population characterized by high infection and/or vaccination rates resulted in immune evasion and distinct clinical disease. Our analysis showed that the epitope profiles of each variant explained the respective potential for cytokine production, including the variants that were more likely to cause cytokine storms. Finally, it serves as a mean to explain the multi-wave patterns observed during SARS-CoV-2 pandemics.

## 1. Introduction

The first cases of COVID-19, caused by the Severe Acute Respiratory Syndrome Coronavirus 2 (SARS-CoV-2), were reported in December 2019 in Wuhan City, China. The virus rapidly spread worldwide, reaching Brazil on February 26, 2020 (source: https://covid.saude.gov.br), when SARS-CoV-2 was detected in a 61-year-old male traveler returning from Italy. As of June 2023, SARS-CoV-2 has infected over 37,656,050 individuals in Brazil with 1.8% case fatality rate and all Brazilian states had documented disease (1).

SARS-CoV-2 is the seventh coronavirus in a diverse family of enveloped positive-sense single-stranded RNA viruses known to infect humans. It has a genome of 29,903 kb and most of its genome codes for ORF1ab (∼72%) which is involved in viral replication and pathogenesis, while other ORFs code for structural (Spike [S], envelope [E], membrane glycoprotein [M] and nucleocapsid [N]) and accessory proteins (2). The spike protein is a target of study since its critical ability to mediate viral cell entry. Once the virus interacts with the host cell, the spike protein undergoes structural rearrangement, enabling the virus to fuse with the host cell membrane (3). Overall, the structure of the spike protein consists of an extracellular N-terminus, a transmembrane domain anchored in the viral membrane, and a short intracellular C-terminal segment (4). Its S1 subunit contains a signal peptide located at the N-terminus and receptor binding domain (5). The S2 subunit comprises the fusion peptide (FP), heptapeptide repeat sequence 1 (HR1), heptapeptide repeat sequence 2 (HR2), transmembrane domain and cytoplasm domain (6).

The emergence of mutations through successive virus replication, a natural feature of RNA virus, led to new SARS-CoV-2 variants and new challenges for curbing the transmission and disease, in spite of massive vaccination coverage in Europe and Americas since January 2021. The majority of the mutations accumulated in the spike protein, and such mutations have been linked to altered efficiency in S1-S2 cleavage which is required for SARS-CoV-2 cell entry, increased tropism and pathogenicity, and virus transmissibility (7–10). The spike protein, and more specifically the receptor binding domain (RBD), is currently the target for the development of COVID-19 neutralizing antibodies and new vaccines (11, 12).

Intriguingly, SARS-CoV-2 variants had a significant impact on the number of cases and deaths in different parts of the globe, as seen when the Gamma variant emerged in South America (13–15) and the Delta and its sublineages in other parts of the globe (16, 17). Those variants had distinct pathogenesis because of immune escaping from prior immunity generated through natural infection and/or vaccination (18, 19).

From March 2020 to December 2022, 5 variants circulated in Brazil (Zeta, Gamma, Alfa, Delta, and Omicron), causing changes in the dynamics of confirmed cases and deaths. The VOC Gamma (P.1) was first identified in early December 2020 in Manaus, Amazonas State, Northern Brazil, and was associated with a higher virus transmissibility (20) which quickly spread across the country (21) and was detected in the state of Rio Grande do Norte early January 2021 (22). In Brazil, the predominance of the Gamma variant was marked by an increased number of new daily cases and deaths (23). Then, by late April, the Delta variant (B.1.617 and AY.*) was introduced (24, 25) and replaced the Gamma variant cases in July 2021 (23), characterized by a low number of new daily reported cases (26) with apparent less fatality, although in the United States of America the Delta variant was associated with a higher number of cases (27). In January 2022, the number of cases increased abruptly again due to the introduction of the Omicron (BA.*) variant; however, its lethality did not follow the proportion of people infected in Brazil and other countries (26).

Most viral infections require the adaptive immune system to be controlled. The strategic capabilities of B cells, CD4+ T cells, and CD8+ T cells present different roles in viral infections and responses to vaccines, it is critical to study adaptive immunity to SARS-CoV-2. In a cellular adaptive response, SARS-CoV-2 infection elicits CD4+ and CD8+ T cells that are directed against a variety of antigens, including structural and nonstructural proteins, and are significantly associated with milder disease (28). Agerer and collaborators showed that changes in CD8+ T epitopes interfered with cytokine production and with cytotoxic activity, which can subvert this cell type surveillance ability (29). Neutralizing antibodies can block the binding viral spike protein to the ACE2 receptor and can promote effector function (30) against the virus. Changes in T cell epitopes promote immune dysregulation, including increase of cytokines as TNFα, IL1 and IL6, as well as chemokine (IL8) which resulted in outraged inflammation with a production of cytokine storm and further worsening of the disease as seen within the first 18 months of SARS-Cov-2 circulation (29, 31–35).

The state of Rio Grande do Norte, northeast Brazil, was used as a model to understand the genetic diversity of the SARS-CoV-2 spike protein. The state has 1.7% of the Brazilian population and between March 2020 and July 2022 reported 710,000 COVID cases and more than 8,000 deaths. The SarsCoV2 sequences from the state of Rio Grande was used as a model to scale up the insights of global spread dynamics, using large-scale global efforts to understand and interpret immunological data using bioinformatics approaches of virus sequenced in the locality. The aim of this study was to apply immunoinformatics to investigate the variation in cluster of SARS-CoV-2 variants in a Brazilian state characterized by a high level of population immunity resulting from both natural infection and vaccination.

## 2. Methods

### 2.1. Data acquisition and filtering

COVID19 data was obtained from Rio Grande do Norte State, Brazil, (LAIS, https://covid.lais.ufrn.br/#dados). The RN vaccination data was downloaded from Open Data SUS (https://opendatasus.saude.gov.br/dataset/covid-19-vacinacao/) up to November 3, 2022. We chose to use only the first and second doses, or the single-dose vaccine, for this analysis. Additional booster doses began to be used in July 2021; however, we decided to disregard this information because of the lack of systematized dose nomenclature by OpenData SUS database.

The translated amino acid sequence isolated was obtained from GISAID (36), downloaded on August 9, 2022, and a complete list of identifiers is available in Supplemental Material 1. Ambiguous amino acid residues were removed, and those sequences shorter than 1209 residues, which represents 95% of the 1273 aa from the spike reference, NC_045512, were excluded from the analysis. Then, the remaining sequences were aligned by MAFFT (37) with the *--add* option, using the GISAID spike alignment from the NC_045512 sequence as seed. The data acquisition, clearing, and the following steps are summarized at Fig. S1 (Supplemental Material 2).

### 2.2. Protein clustering

The CD-HIT performed the protein clustering analysis (38). To determine the optimal clustering threshold, GISAID-selected variant spike alignment was used. The clustering analysis began with a cut-off of 100% identity and reduced by 0.1% until it reached 98.0% (Fig. S2). A 99.0% limit was used since CD-HIT was able to cluster the lineages of wild type (B.1.*), Gamma (P.1*), and Omicron (BA.*) into different clusters which were compatible with dynamics of infection. Following the previously established cut-off point, the retrieved spike sequences were aggregated using CD-HIT to provide a representative sequence for each cluster, which was then employed as an input for epitope prediction algorithms. The order of the clusters was kept the same as the CD-HIT output.

### 2.3. Epitopes identification

#### 2.3.1. Linear CD4+ T cell epitopes prediction

For MHC-II-binding epitopes, 15-mer long epitopes for each cluster from spike protein were predicted using NetMHCII 2.3 (39) server and confirmed with NetMHCIIpan 4.0 (40) server. We selected for T-cell epitope prediction the HLA class II molecules *DRB1*0101, DRB1*0301, DRB1*0401, DRB1*0701, DRB1*0802, DRB1*0901, DRB1*1101, DRB1*1302, DRB1*1402, DRB1*1501, and DRB1*1601*, which are most prevalent in Latin America (41). The predicted peptides were classified as strong and weak binders with a threshold percentile rank (% Rank) ≤ 2% and ≤ 10%, respectively. The non-binder peptides (> 10% rank) were not considered in the study.

#### 2.3.2. Linear CD8+ T-cell epitopes prediction

The prediction of peptides (9 amino acids long) in linear form to the MHC class I groove was performed by NetMHCpan 4.1 (40) that applies an Artificial Neural Networks (ANNs) method. The threshold for binding affinity of peptide-MHC-I (percentile rank) was set at 0.5% for strong binders and 2% for weak binders. Additionally, NetCTL (42) was used to confirm predicted epitopes. This web server requires the protein sequence in its FASTA format in order to carry out various analyses such as MHC class I binding affinity prediction, the efficiency of TAP transport, and C-terminal cleavage activity. The HLA alleles selected for the study included: *HLA-A01:01, HLA-A02:01, HLA-A02:04, HLA-A02:11, HLA-A02:19, HLA A24:02, HLA-A24:03, HLA-A26:01, HLA-A31:01, HLA-A68:01, HLA-B07:02, HLA B08:01, HLA-B15:01, HLA-B15:04, HLA-B15:07, HLA-B27:05, HLA-B35:01, HLA B35:04, HLA-B35:05, HLA-B35:11, HLA-B35:18, HLA-B35:19, HLA-B39:01, HLA B39:03, HLA-B39:05, HLA-B39:06, HLA-B39:09, HLA-B40:02, HLA-B40:04, HLA B48:01, HLA-B48:03, HLA-B51:01, HLA-B51:04, HLA-B52:01,* and *HLA-B58:01*, which are most prevalent in Latin America (41).

#### 2.3.3. Linear B cell epitopes prediction

For prediction of linear B epitopes of spike Sars-Cov2, the sequences were submitted to the online tool IEDB (Immune Epitope Database) (http://tools.immuneepitope.org/bcell/), using the method of Kolaskar and Tongaonkar (43). The method is based on the occurrence of amino acid residues in experimentally determined epitopes. Application of this method can predict antigenic determinants with about 75% accuracy (43). To confirm the epitopes obtained, the BCEpred web server (http://crdd.osdd.net/raghava/bcepred/) was used. This web server is based on a plethora of physicochemical propensity scales utilizing amino acid properties, such as hydrophilicity and antigenicity, either individually or in combination. Antigenic propensity (Kolaskar) was selected, and a threshold of 1.9 was used. Results from the two prediction tools were aligned simultaneously, and overlapping sequences were assumed to be B-cell epitopes.

#### 2.3.4. Conformational B epitopes

ElliPro (44) is an online bioinformatics web server that has been used for the prediction of conformational epitopes based on the 3D structure of spike proteins from each cluster. The web server uses three algorithms: approximation of a protein surface patch by an ellipsoid, computation of the residue protrusion index (PI), and clustering neighboring residues based on their PI values. The threshold was fixed at 0.5.

#### 2.3.5. Protein modelling

The SARS-CoV-2 spike (S) proteins from each cluster were applied to the I-TASSER server (45). For each target, I-TASSER simulations generate a large ensemble of structural conformations called decoys. To select the final models, I-TASSER uses the SPICKER program to cluster all the decoys based on pair-wise structure similarity and reports up to five models corresponding to the five most significant structure clusters. The confidence of each model is quantitatively measured by the C-score, which is calculated based on the significance of threading template alignments and the convergence parameters of the structure assembly simulations. Later, based on the TM-score, the TM-align algorithm was used to identify the best model structures (46).

### 2.4. Prediction of cytokine-stimulating ability

#### 2.4.1. IL-10 Pred and IFN epitope analysis

Epitopes inducing IL-10 were identified as described by (47). To assess their ability to induce IFN-gamma and IL-10 cytokines, the unique epitopes from each cluster of SARS-Cov2 were submitted to the IFNepitope (DHANDA; VIR; RAGHAVA, 2013) and IL10 Pred (NAGPAL et al., 2017) web servers. The IFNepitope Server predicts IFN-inducing epitopes from a set of peptide sequences. As a result, for each cluster, this server was used to predict IFN-inducing unique epitopes. The prediction search model was an SVM (support vector machine) motif hybrid, which provides results by combining the SVM-based model with motifs that contribute to IFN-induction. The model performs better compared with an SVM model and a motif-based model separately. Similarly, the IL-10 Pred web server was used, which allows users to discover, design, and map peptides with the capability to induce the IL-10 cytokine. The IL-10 inducing ability of unique epitopes from each cluster was determined using the IL-10 Pred with an SVM-based prediction model. In the case of IL-10 Pred, we used the default cutoff values as recommended by developers.

#### 2.4.2. Predicting IL-6 inducing epitopes

The unique epitopes from each cluster of SARS-Cov2 spike protein were subjected to IL-6-Pred (48). To predict the secretion of IL-6, the IL-6-Pred uses a wide range of machine learning techniques, out of which a random forest-based model achieves the maximum accuracy on the dataset evaluated. To consider a peptide an IL-6 inducer, the default value of > 0.11 is recommended.

#### 2.4.3. Predicting IL-4 inducing epitopes

The unique peptides from each cluster were subjected to the IL-4-Pred (49) server, which predicts the ability of these peptides to induce Interleukin-4 (IL-4). The IL-4-Pred is based on SVM (Support Vector Machine-based Methods) and the motif algorithm, which incorporates the amino acid composition and propensity, di-peptide composition, and physico-chemical properties of a peptide for prediction. To consider a peptide an IL-4 inducer, the default value of > 0.2 is recommended.

#### 2.4.4. Pro-inflammatory peptides

The unique epitopes from each cluster of SARS-CoV-2 S protein were subjected to the PIP-EL server (50) to predict pro-inflammatory responses. In PIP-EL, a peptide was considered pro-inflammatory if it could induce cytokines like IL1α, IL1β, TNFα, IL6, IL8, IL12, IL17, IL18, and IL23. The PIP-EL server is meant to be a cumulative dataset with ten independent models for amino acid sequence, di-peptide composition, distribution-transition, and physicochemical properties. Any score value greater than 0.45 considers a peptide to be an inducer of a pro-inflammatory cytokine.

#### 2.4.5. Anti-inflammatory peptides

The unique epitopes from each cluster of the SARS-CoV-2 spike protein were subjected to the Pre-AIP (Prediction of Anti-Inflammatory Peptides) server (51), and these peptides were evaluated for being anti-inflammatory in nature. A peptide was considered anti-inflammatory if it could induce anti-inflammatory cytokines like IL-10, IL-4, IL-13, IL-22, TGF-b, IFN-⍺ and IFN-β. Pre-AIP systematically investigates different types of characters, which include primary sequence, evolutionary, and structural information, through a random forest classifier (51). A high-confidence AIP was labeled as having a score range of >0.468, 0.468 > score ≥ 0.388 was labeled as a medium-confidence AIP, and 0.388 > score > 0.342 is labeled as low confidence.

## Data availability

The sequences of SARS-CoV-2 spike glycoprotein analyzed in this study were downloaded from the Global Initiative on Sharing All Influenza Data (GISAID).

## 3. RESULTS

### 3.1. COVID-19 Epidemiology and vaccination rates in Rio Grande do Norte state

The COVID-19 pandemic scenario in Brazil and elsewhere was characterized by multiple waves of virus dissemination, resulting in increasing numbers of cases and deaths, with peaks followed by valleys (52). Between March 2020 and June 2022, a total of 579,931 confirmed COVID-19 cases, this number is probably underrepresented since there was not availability of tests for all symptomatic subjects in the first 12 months of the pandemic, and 8,315 deaths were reported in the state of Rio Grande do Norte, Brazil, representing a case fatality rate of 1.4%. During this period, the first wave occurred between May and September 2020, accounting for 88,147 cases (15.2% of the total) and 2,477 deaths (30.1% of the total), with a case fatality rate of 2.8%. The second wave spanned from February to August 2021, contributing 193,054 cases (33.3% of the total) and 3,838 deaths (46.1% of the total), resulting in a case fatality rate of 1.9%. The third wave, which took place from January to March 2022, accounted for 161,995 cases (27.9% of the total) and 501 deaths (6.0% of the total), with a case fatality rate of 0.03% (Fig. 1A).

**Figure 1.**
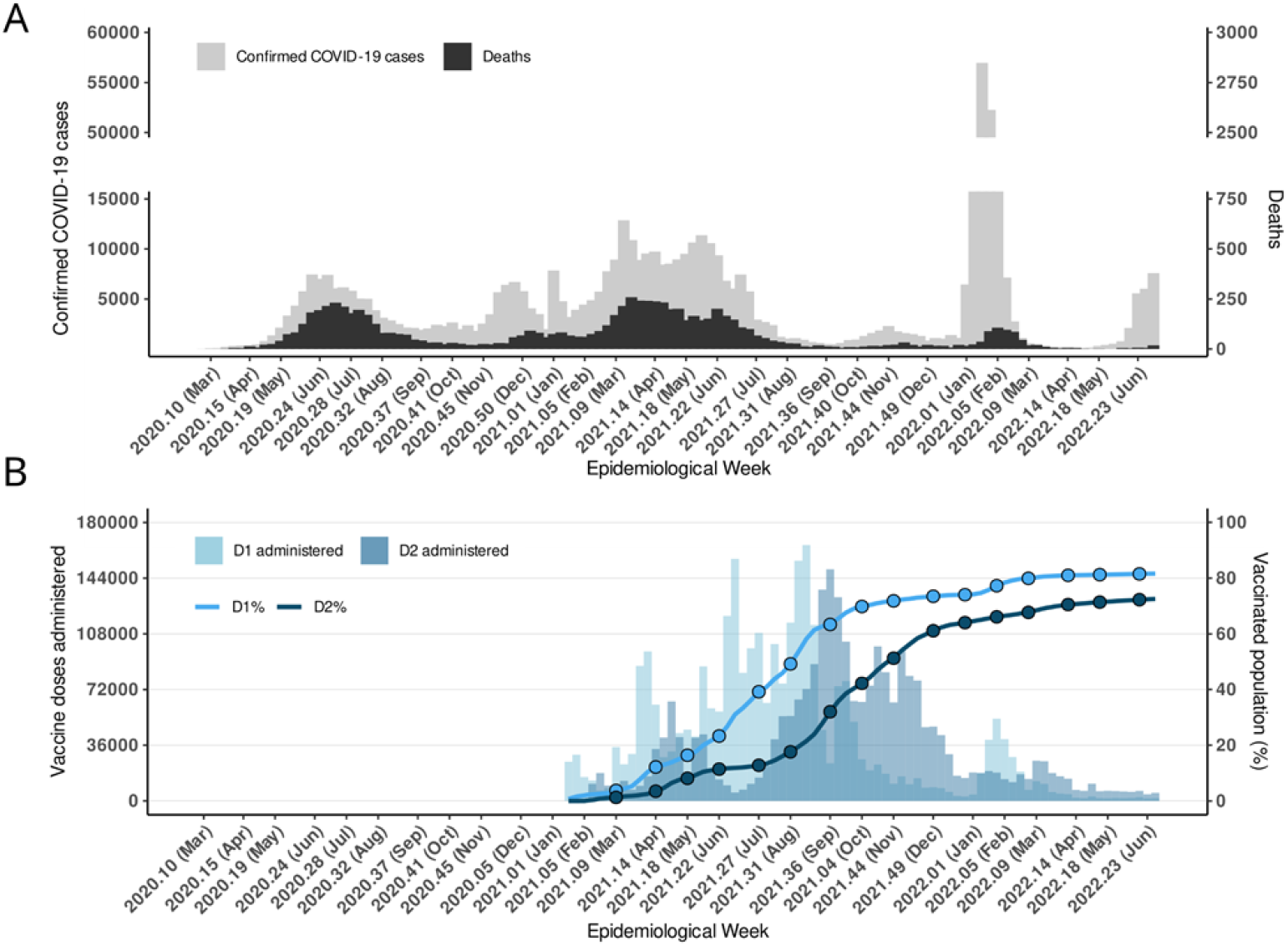
COVID-19 reports in Rio Grande do Norte State, Brazil. (A) Confirmed cases and deaths of COVID-19 reported by the Public Health Department of Rio Grande do Norte State - SESAP/RN. (B) COVID-19 Vaccination status obtained from OpenData SUS.

The immunization programs started on January 19, 2021 (26), and by August 2021, approximately 40% of the population had received at least one shot (D1). By the end of 2021, 74% of the population had received the D1 shot, while 64% had received a booster (Fig. 1B). As of mid-June 2022, 81.6% and 72.5% of the population had received D1 and D2, respectively.

### 3.2. The evolution of SARS-CoV-2 variants in the Rio Grande do Norte state, Brazil

Using raw SARS-CoV-2 genomic data from GISAID (Fig. 2A), from 1715 sequences, at least 5 lineages were detected in the RN state, in chronological order: the Wild-type; Zeta (P.2); Gamma (P.1 and P.1*); Alfa (B.1.1.7 and Q.*); Delta (B.1.617.2 and AY.*); and Omicron (B.1.1.529 and BA*). In November 2020, the Zeta variant was discovered co-circulating with non-VoCs or VoCs isolates. The Gamma lineage emerged in January 2021 and quickly rose to dominance by late February, persisting until August 2021. Between April and late July 2021, the Delta variant was detected sporadically and at a low frequency, but it became dominant from August to December 2021. The Omicron variant was initially detected in November 2021; however, it rapidly emerged as the prevailing lineage by January 2022, and its subsequent subvariants successively gained prominence until the conclusion of this temporal analysis as of June 2022.

**Figure 2.**
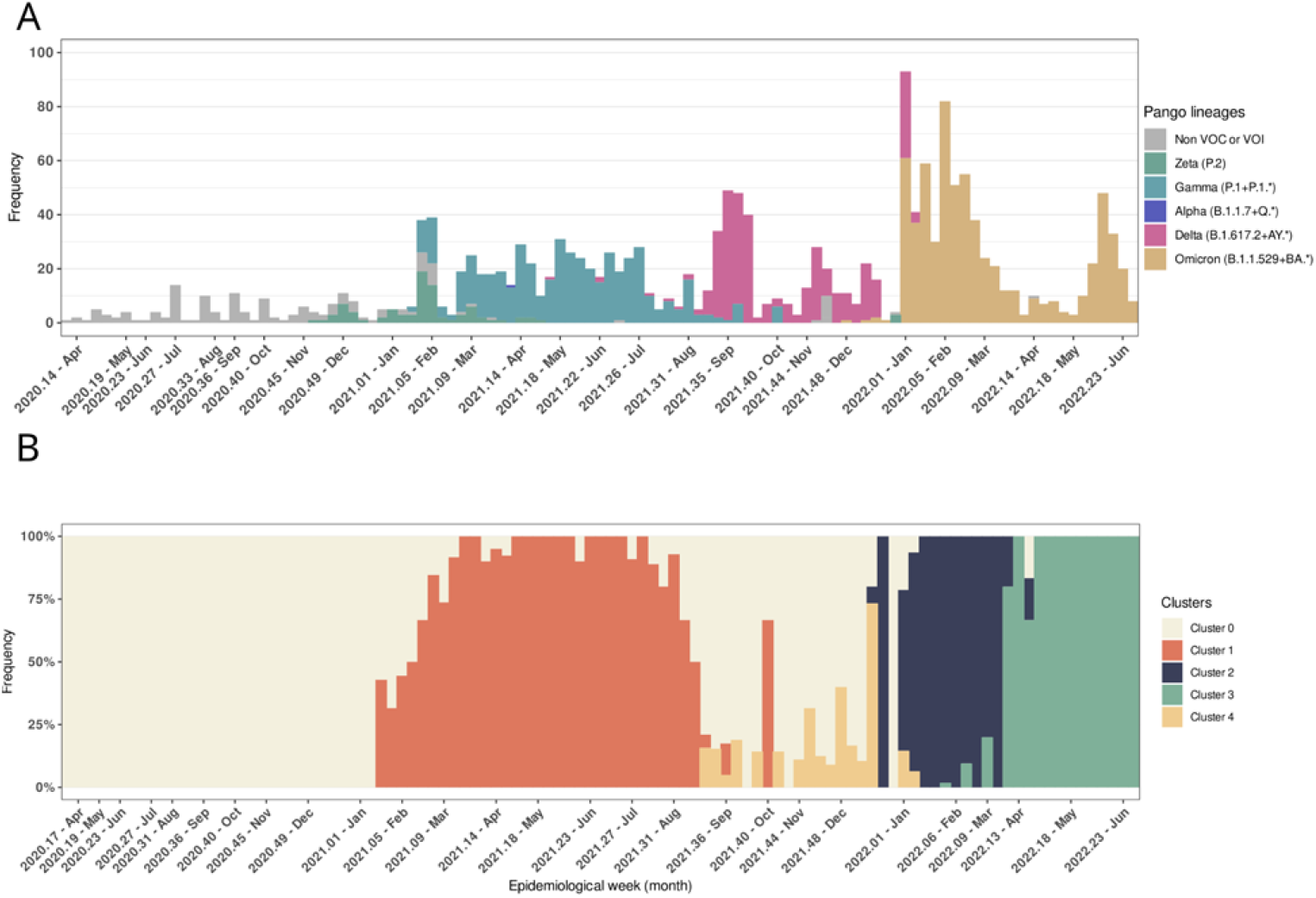
SARS-CoV-2 genomic features in the Rio Grande do Norte State. (A) SARS-CoV-2 lineages circulated throughout the pandemic period based on the genetic sequences deposited in GISAID. (B) Clusters composed of Spike proteins are distributed based on the isolation period of the variant. The cluster names have been preserved in the same format as defined by CD-HIT.

### 3.3. Five clusters of SARS-CoV-2 spike protein sequences were identified between March 2020 and July 2022

The spike protein sequences were then grouped based on their similarity. The original data set containing 1715 amino acid sequences were filtered to ensure the quality for the input for cluster analysis, with removal of ambiguous residues and exclusion of sequences shorter than 1209 aa, resulting in 1257 sequences (Fig. S1). Using 99.0% identity as a cut-off for cluster analysis, we observed five groups named Cluster 0, Cluster 1, Cluster 2, Cluster 3, and Cluster 4 among the spike protein sequences (Fig. 2B). Of importance, the cluster numbers were conserved in the format originally defined by CD-HIT and do not necessarily reflect the chronological order in which the sequences within each cluster emerged in the region. Cluster 0 comprises lineages that emerged predominantly between March 2020 and January 2021 (the wild type strain and the Zeta variant) and then between mid-August and December 2021 (some Delta lineages). Cluster 1 is entirely composed of Gamma lineages and showed a predominance from February to August 2021. The Delta lineages were distributed across Clusters 0 and 4 as observed between August and December 2021. Indeed, Cluster 4 was entirely composed of Delta lineages. Finally, Clusters 2 and 3 contained Omicron variants that emerged in late December 2021. SARS-CoV-2 predicted epitopes revealed likely COVID-19 occurrence patterns

#### 3.3.1. Prediction of linear epitopes

To investigate the linear B-cell epitopes in the five distinct clusters of SARS-CoV-2 spike protein, we combined predictions from two immunoinformatics approaches, Bepipred 2.0 and IEDB (Kolaskar & Tongaonkar Antigenicity). We observed that, with the exception of Cluster 1, which presented 27 linear epitopes, we identified 28 B-cell linear epitopes in each cluster (Supplemental Material 3) and 14 epitopes shared among all five B-cell clusters (Fig. 3A). The CD4+ T cell linear epitopes were predicted by the combination of NetMHCII v2.3 and NetMHCIIpan v4.0 softwares, and resulted in 81, 64, 67, 70, and 62 CD4+ T-cell epitopes for Clusters 0, 1, 2, 3 and 4, respectively (Supplemental Material 4), depicting 51 shared epitopes among all five clusters (Fig. 3B). For CD8+ T-cell epitopes, we observed 149, 155, 151, 148, and 152, respectively for Cluster 0, 1 2, 3 and 4 (Supplemental Material 5). All five clusters shared 117 CD8+ T linear epitopes identified by combining NetMHCpan v4.1 and NetCTL v1.2 (Fig. 3C).

**Figure 3.**
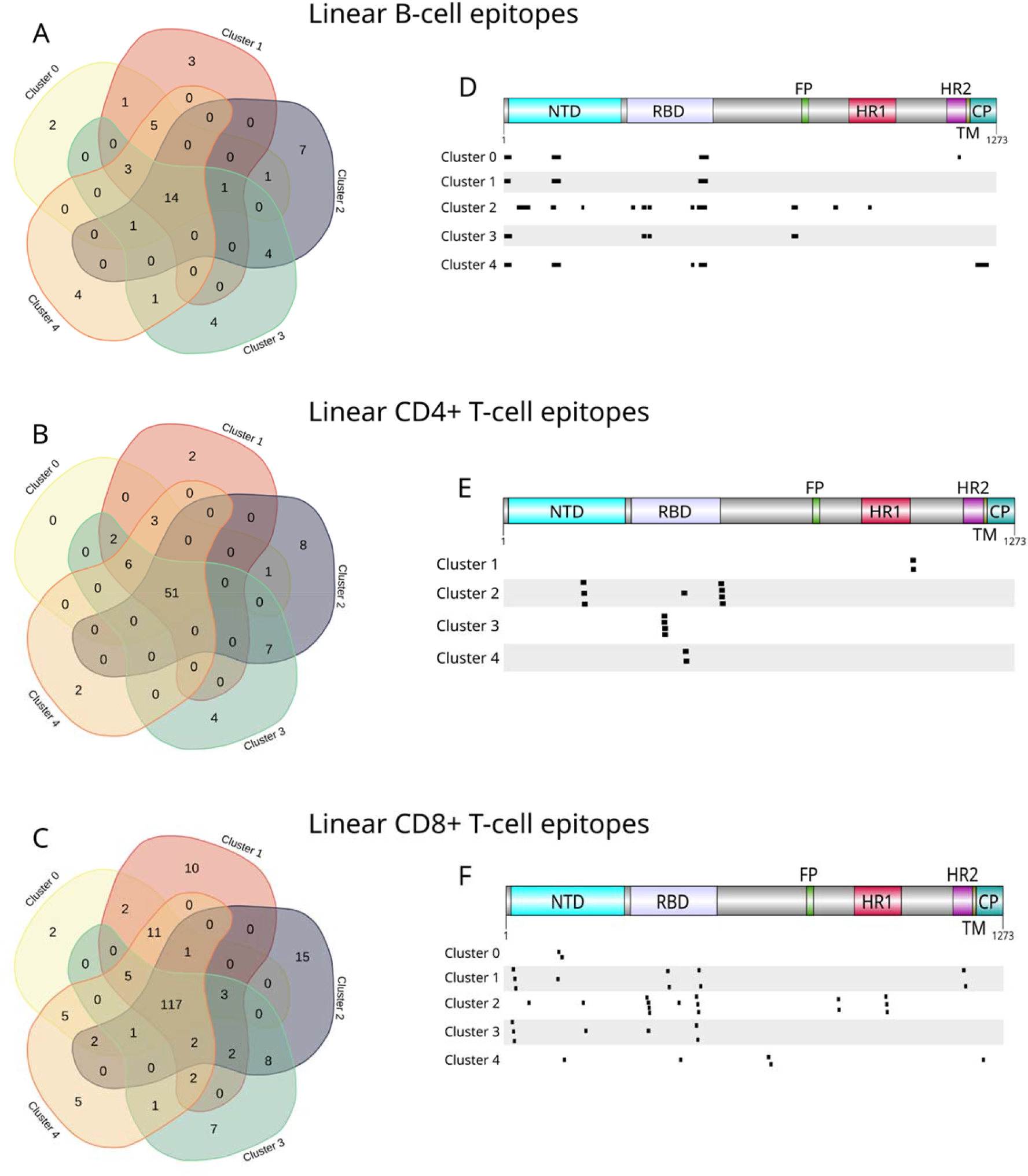
Predicted linear epitopes. (A, B, and C) Venn diagram illustrates identical linear B cells, CD4+ T cells, and CD8+ T cells cell epitopes among the five clusters. (D, E, and F) The scheme depicts the position of each unique predicted linear epitope over the spike protein primary sequence for B cells, CD4+ T cells, and CD8+ T cells in all clusters.

We focused on the distinct features among the clusters to understand the dynamics of COVID19 as the virus evolved. Through the analysis of these clusters, we examined distinct epitopes that may provide insights into the intrinsic factors associated with the outcomes from variant. As a result, Cluster 0 had the fewest unique epitopes; Cluster 1 showed variability, primarily for CD8+T epitopes; Cluster 2 featured the most distinct epitopes; and Clusters 3 and 4 had a decreasing pattern of unique epitopes. The majority of the unique epitopes for each cluster are mapped into spike protein regions that cover the NTD and RBD domains (Fig. 3D, E, and F).

#### 3.3.2. Conformational B cell epitopes

The Ellipro method was used to predict discontinuous (non-linear) B-cell epitopes based on the 3D structures of each cluster’s representative spike proteins. Although the discontinuous epitopes were predicted independently for each cluster’s model, we chose to unify a single model to show the regions where these epitopes were identified (Fig. 4A). We analyzed conformational B-cell epitopes to assess the changes that could reduce the effectiveness of potential neutralizing antibodies (53). Clusters 0 and 1 displayed similar discontinuous epitopes region with minor changes in the amino acid residues comprising them (Fig. 4B and C; Supplemental Material 6). Clusters 2 and 3 contained more immunogenic residues in non-cytoplasmic domains of the protein (Fig. 4D and E). Cluster 4 showed the greatest diversity of discontinuous epitopes, including the entire region of the transmembrane and cytoplasmic domain as antigenic (Fig. 4F); however, it was not followed by disease severity or deaths (Fig. 1A). Similarly to the linear B epitopes, NTD and RBD figure as important sites for epitope identification in all SARS-CoV-2 lineages that have been widespread in the State of Rio Grande do Norte.

**Figure 4.**
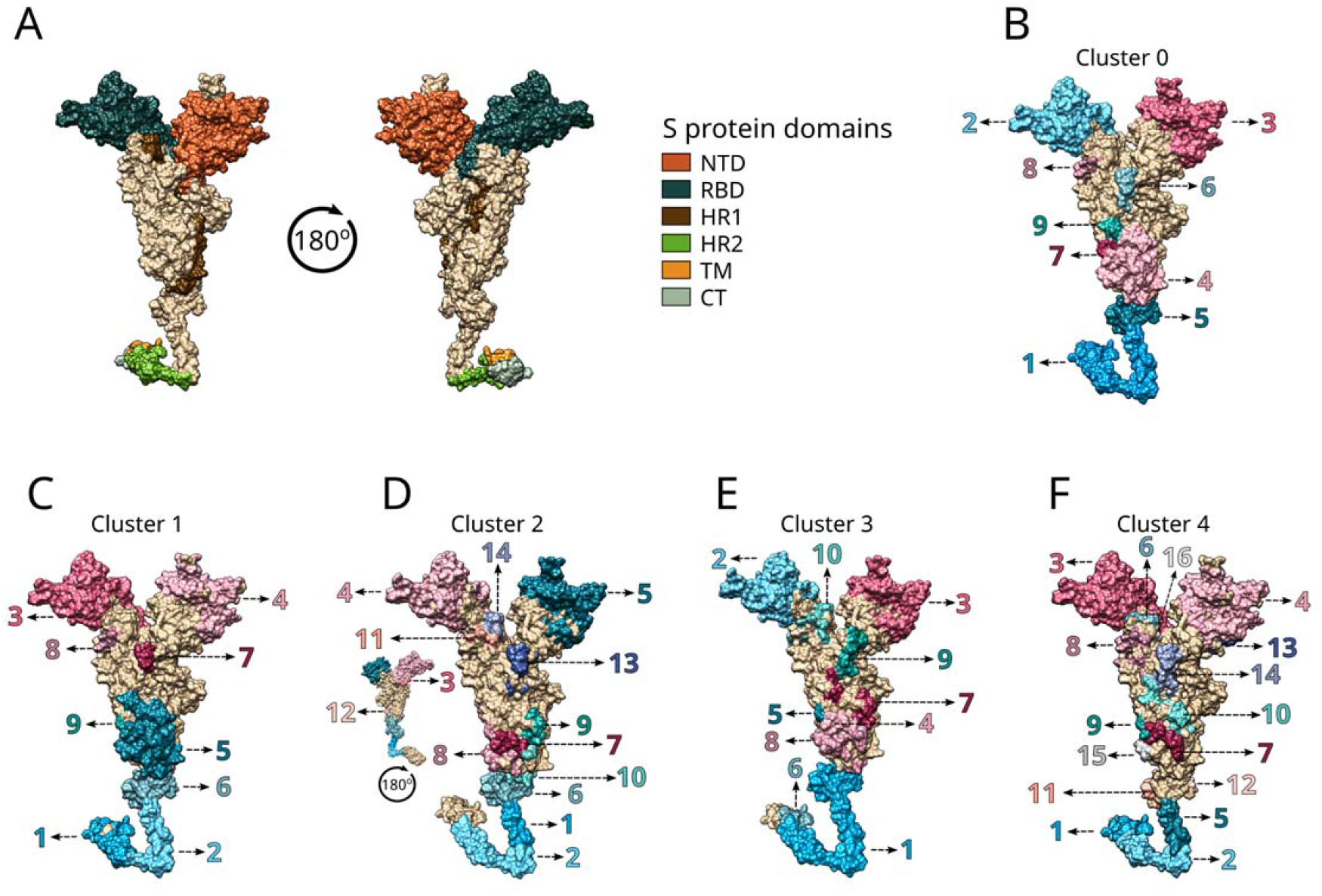
Predicted conformational B-cell epitopes. (A) Schematic representation of the SARS-CoV-2 S protein indicating the position of the domains NTD (N Terminal Domain); RBD (Receptor-Binding Domain); HR1 (Heptad Repeat 1); HR2 (Heptad Repeat 2); TM (Transmembrane Domain); and CT (Cytoplasmic Tail Domain) over the predicted tertiary structure. (B, C, D, E, F) Predicted conformational B-cell epitopes respectively for clusters 0, 1, 2, 3 and 4. The different colors and the numbering, indicating the position of the epitope, are defined by the Ellipro output order.

### 3.4. Prediction of cytokine production *in silico* by unique T and B linear epitopes

Cytokine dysregulation which led to the so-called cytokine storm which were majorly reported within the first 18 months of SarsCov2 circulation resulted in increased morbidity and mortality (54). We evaluated the influence of the predicted epitopes over the induction of IFN-gamma, IL-6, IL-4, and IL-10. We also evaluated the overall induction of pro-inflammatory cytokines (IL1α, IL1β, TNFα, IL6, IL8, IL12, IL17, IL18, and IL23) and the capacity of the epitope to induce an anti-inflammatory response, as IL-10, IL-4, IL-13, IL-22, TGF-beta, IFN-alpha, or IFN-beta. As a result, rather than analyzing cytokines individually, we considered whether the set of the cytokines had either pro- and anti-inflammatory activities.

#### 3.4.1. Cluster 0

Following analysis, CD4+ T cells were not predicted to have any unique epitopes in this cluster (Table 1). The predicted CD8+ T cell epitopes have the potential to elicit both anti-inflammatory and pro-inflammatory responses (Table 1). A similar result was observed for the unique B cell epitopes (Table 1).

**Table 1.**
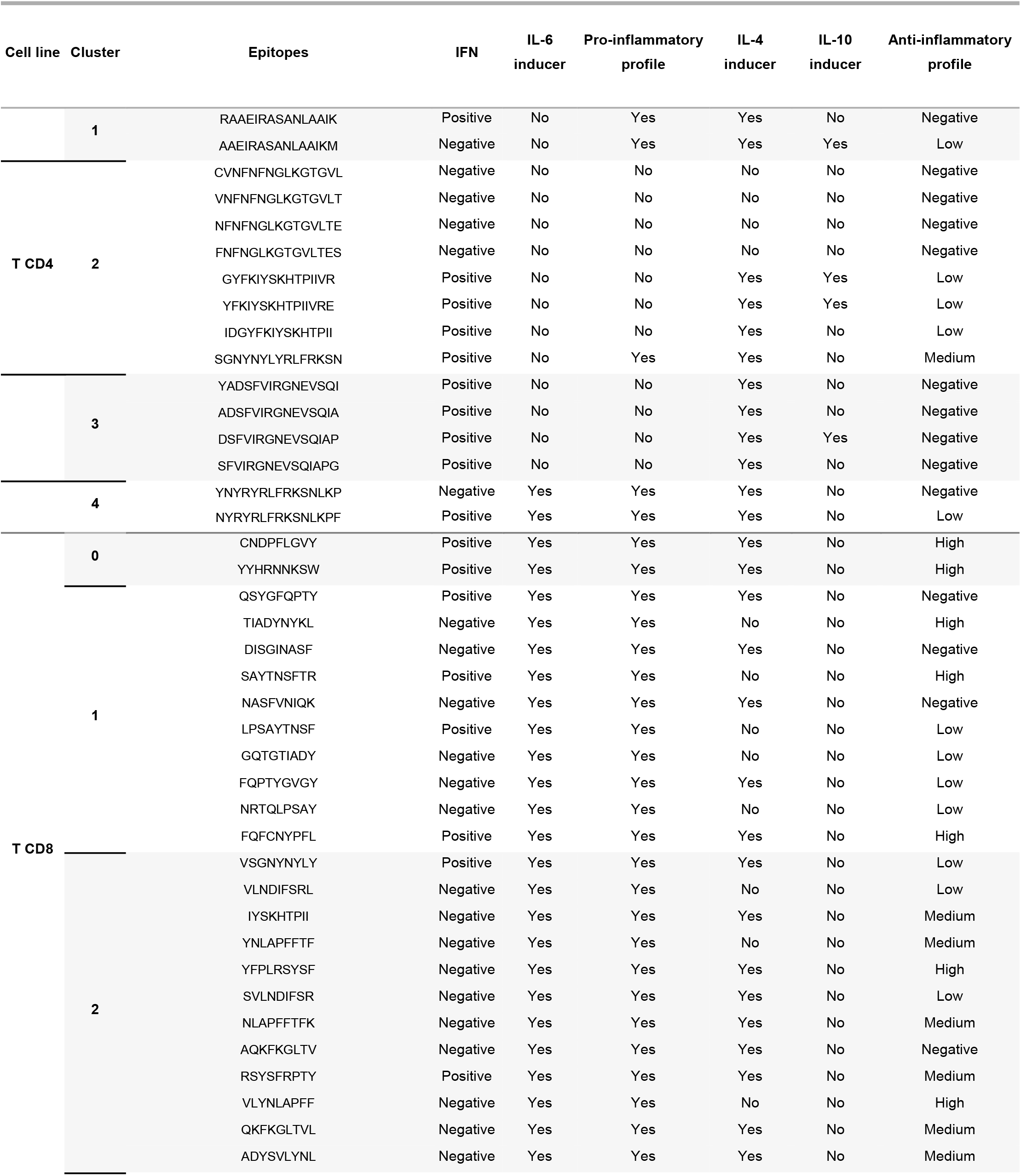
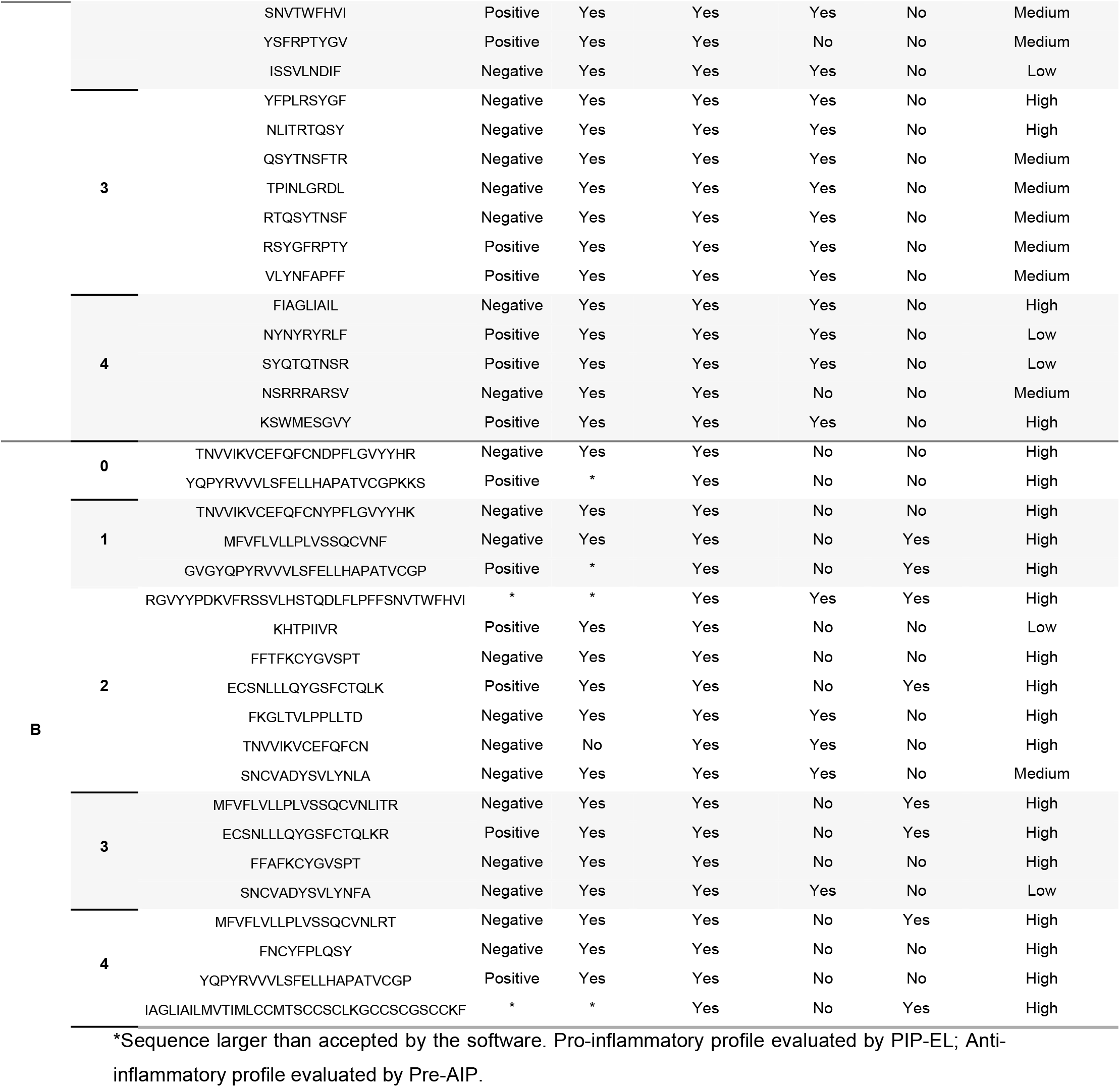
Unique linear CD4+ T, CD8+ T, and B cell epitopes cytokine profile induction for each cluster.

#### 3.4.2. Cluster 1

Two unique CD4+ T cell predicted epitopes display the potential ability to elicit pro-inflammatory responses with a low or no anti-inflammatory profile; however, they were able to induce IL-4 and IL-10. All predicted CD8+ T cell epitopes have the potential to elicit pro-inflammatory responses (100% are potential IL-6 inducers) and the majority had a low or no anti-inflammatory properties induction (8/10 epitopes). Concerning the predicted B cell epitopes, all of them showed potential pro-inflammatory (the majority are potential IL-6 inducers) and anti-inflammatory responses (the majority are potential IL-10 inducers).

##### Cluster 2

Only one of the eight unique CD4+ T cell predicted epitopes evoked pro-inflammatory responses, and the chances of eliciting anti-inflammatory responses were poor. In contrast, all predicted CD8+ T cell epitopes had the potential to elicit pro-inflammatory responses (100% are potential IL-6 inducers), but the majority had the ability to induce anti-inflammatory responses as well (11/15 [73.3%] are IL-4 inducers). Regarding the B cell epitopes predicted, they all had pro-inflammatory characteristics (5/6 [83.3%] and were IL-6 inducers) and also displayed an anti-inflammatory profile (4/7 [57.1%] are potential IL-4 inducers).

##### Cluster 3

Although the CD4+ T cell predicted epitopes do not display a pro-inflammatory profile and show 100% IL-6 non-inducers, they are INF-gamma inducers and they did not show anti-inflammatory potential responses, even though they were IL-4 inducers. In addition, all predicted CD8+ T and B cell epitopes had the potential to elicit pro-inflammatory responses and were 100% potential IL-6 inducers. For predicted CD8+ T cell epitopes, they may induce a high- or medium-anti-inflammatory profile and 100% IL-4 inducers. Instead, for the B-cell epitopes predicted, the majority were IL-4 non-inducers; however, most of them displayed an anti-inflammatory profile as well.

#### 3.4.3. Cluster 4

Similarly as observed with previous clusters, the CD4+ T, CD8+ T, and B cell predicted epitopes display potential to elicit pro-inflammatory responses, and the majority are anti-inflammatory inducers, with the exception of CD4+ T cell predicted epitopes in which the same pattern was not seen for the anti-inflammatory profile.

## 4. Discussion

During COVID-19 the first three years of the COVID19 pandemic, the SARS-CoV-2 mutated and evolved with genetic variation in the population of circulating viral strains and after over 48 months of circulation within the human population it continues to mutate. The emergence of more recent SARS-CoV-2 variants, characterized by heightened transmissibility, raises concerns regarding their potential to compromise the protective effects of adaptive immunity elicited through natural infection or vaccination, since apparently immunity to either cases may be short lived (19), as we have observed here previously. Nevertheless, it is important to note that previous immunity, whether acquired through prior infection or vaccination, continues to demonstrate its association with protection against COVID-19 severity (55).

More than 739,000 people in the northeastern Brazilian state of Rio Grande do Norte were confirmed to haver COVID-19 in the previous two years; although one needs to consider that in the first 12 months of the pandemic there was not readily availability of RT-PCR for all symptomatic subjects, therefore this total is underrepresents the total number of infected subjects. There were nearly 9,000 deaths (26). COVID-19 progressed in this state along the same timeline as the rest of Brazil, presenting three distinct waves that were observed associated with novel variants.

Our cluster analysis of the virus sequences was conducted to explore whether factors such as prior infection, vaccination, the prevalence of less virulent strains, or a combination of at least two of these factors, were associated with the estimated potential of inflammatory and anti-inflammatory epitopes and, consequently, clinical outcomes.

We identified *in silico* linear and discontinuous peptides representative of B- and T-cell epitopes that were conserved and unique across the SARS-CoV-2 spike glycoprotein sequences obtained from each SARS-COV-2 variant that circulated in the state of Rio Grande do Norte State between March 2020 and July 2022. Nonetheless, we focused on the unique epitopes for each cluster in order to highlight possible explanations for the temporal increase and decrease in the number of cases and deaths caused by COVID-19. We defined five clusters of SARS-CoV-2 lineages by using a clustering method on 1257 high-quality spike protein sequences.

In early pandemics in the state of Rio Grande do Norte, non-VoCs and Zeta lineages in Cluster 0 caused rapid transmission and high fatality rates, declining by August 2020. The Gamma variant (Cluster 1) with concerning spike protein mutations, which included E484K, N501Y, and K417T, showed increased transmission and severity (56). Cluster 1 introduced three new B-cell epitopes, two CD4+ T-cell epitopes, and sixteen CD8+ T-cell epitopes.

Further, the Delta variant (lineages included in Cluster 0 and Cluster 4) became prevalent from mid-August 2021 to late-December 2021, with the lowest daily number of new cases since the start of the COVID-19 pandemic. This trend was observed in various parts of the country, with slight variations in timing and the southeast region being at the forefront of each wave (57, 58). Chronologically, SARS-CoV-2 Cluster 4 lineages were identified in the state of Rio Grande do Norte before the Omicron variant upsurge (sequences included in Clusters 2 and 3). Cluster 4 is composed of a restricted number of Delta lineage spike sequences, which underwent an evolutionary process that led to the emergence of three clades and over 240 sublineages in less than a year (25). Despite the spike sequence differences between Clusters 1 and 4, Gamma and Delta variants co-circulated in the region from August to December 2021; however, the frequency of Cluster 1 sequences was much lower than that of Cluster 4. The prevalence of the Delta variant, during this period, caused no substantial increase in mortality rate in Brazil. Overall, this phenomenon could be linked to the number of individuals with prior immunity either by natural infection and-or by vaccination at that time, which likely limited the upsurge of COVID-19 cases in Brazil due to Delta variant (59).

Our findings revealed that CD8+ T cell epitope variability was higher in cluster 1 than in cluster 4, particularly in the NTD and RBD regions (figure 4F), may explain in part the dramatic decrease of COVID-19 cases and deaths in Brazil at that time. T cell memory encompasses broad recognition of viral proteins and single point mutations can indeed abolish functional responses (60). Additionally, the preservation of the spike protein epitopes was 92% for class II and 95% for class I (61). Although the antibody recognition of the spike protein was reduced for some variants, the decrease for the Delta lineage was moderate (61–63). PLANAS and collaborators (2021) demonstrated that Delta was relatively poorly neutralized by convalescent sera obtained from unvaccinated individuals infected with non-VOC virus; however, when tested against sera from vaccinated individuals, the neutralizing titers were strongly increased. Although the emergence of the Delta variant had a brief journey in Brazil, many countries were heavily affected by the Delta variant, having an outburst of COVID-19 cases and deaths during this wave (60). Also, it is important to point out that these countries were barely affected by the Gamma variant, and they probably did not have new epitopes during the Delta variant wave, which may have led to disease severity and deaths.

Cluster 2 was entirely composed of the Omicron variant (BA.1 and BA.1* lineages), which was detected in late December 2021 and became dominant in January 2022. The Omicron variant introduced a large number of mutations within spike protein, promoting the emergence of new epitopes, primarily in the NTD and RBD regions, with Cluster 2 providing the most distinct unique epitopes across all clusters. Cluster 3 had emerged as the dominant strain by March 2022, consisting of BA.2, BA.2.*, BA.4, BA.4.*, and BA.5 sublineages from the original Omicron variant. Despite its ability to cause immune escape from vaccines (19) or natural infection (65, 66), resulting in a sudden increase in the number of infections, the period following the introduction of the Omicron variant did not show a dramatic increase in hospitalizations and deaths in the state of Rio Grande do Norte, Brazil (67) or worldwide (68–70) compared to the previous COVID-19 waves.

A recent study have shown that changes found in the unique epitopes of each cluster have the potential to bring profound changes in key steps of the immune response, such as reducing or even inhibiting antigen presentation by MHC (29), inhibiting or inducing the production of certain anti- or pro-inflammatory cytokines. It has been recognized that CD8+ T cells play a crucial role in regulating the spread of the SARS-CoV-2 and impacting the intensity of COVID-19 (29). In our study, we observed that a higher number of unique epitopes were identified in CD8+ T cells, in all clusters, compared to CD4+ T and B cells (figure 4). This data reinforces the role of CD8+ T cells in the SARS-CoV-2 infection (71), suggesting that the highest selective pressure occurred in these cells, which may lead to the emergence of VoCs with new epitopes, promoting immune escape. Even vaccinated or previously infected individuals were susceptible to infection and disease development, although with less severity (19). According to Fig. 3 (D, E, and F), the spike RBD contained a substantial amount of unique epitopes that could directly influence the transmission, tropism, and virulence of the virus in humans, as observed and confirmed by the arrival of new variants (72).

The “cytokine storm,” a term coined because of the significant increase in the levels of pro-inflammatory cytokines, was closely related to severity and death because of COVID-19 (54). This phenomenon was considered one of the major causes of acute respiratory distress syndrome and widespread tissue damage resulting in multiple organ failure (54, 73, 74). In addition, the use of anti-IL6 receptor, such Tocilizumab, reduced the likelihood of progression to the composite outcome of mechanical ventilation or death (75, 76). Thus, we investigated the profile of potential cytokine production for each cluster in order to bring up the differences in the disease worsening across multiple outbreaks of infections in RN State.

The unique epitopes of each cluster were subjected to an *in silico* evaluation of their cytokine-inducing potential. Intriguingly, the epitopes showed distinct profiles for certain clusters, which may support our understanding of how the dynamics of infection carried out for some VOCs in circulating in the study area. The unique CD4+ T and CD8+ T cell epitopes (Table 1) display a predominant ability to elicit pro-inflammatory cytokines, which increases the likelihood of severe disease. This was evident when the P1 variant (Cluster 1) emerged and dramatically increased the number of cases and deaths (Fig. 1).

Cluster 2 exhibited the highest count of distinctive epitopes: eight for CD4+ T cells, fifteen for CD8+ T cells, and seven for B cells. This observation raises concerns regarding the potential severity of cases, as the immune system may lack pre-existing immunological memory to mount an effective response. Nevertheless, it is noteworthy that the emergence of the Omicron variant (Cluster 2) was accompanied by a substantial surge in cases, but this increase was not paralleled by a proportional rise in the number of fatalities (Fig. 1 and 2). This could be explained, in part, by data obtained *in silico*, in which most unique epitopes on CD4+ T cells were incompetent of inducing either anti- or pro-inflammatory responses, whereas the majority of unique epitopes on CD8+ T and B cells were capable of eliciting both anti- and pro-inflammatory responses, resulting in a global neutrality response profile, particularly in cluster 2, which makes COVID-19 cases less severe during this period. It is also worth noting that during the peak of the Omicron variant, 65% of the state population had already been vaccinated with at least two doses, potentially reducing the disease severity, as seen elsewhere (77).

To date, this is the first time that the cluster approach has been used as a guide to understand the impact of VoCs throughout the pandemic in the human immune response enabling a perspective of immunosurveillance for the emergence of new VOCs capable of inducing distinct clinical outcomes. The approach could be used for improvement of vaccines in addition to surveillance.

### Limitations and Conclusions

The main limitations of the current study are the relatively restricted number of sequences deposited at GISAID for Rio Grande do Norte state. In addition, with the availability of rapid antibody tests for the detection of SARS-CoV-2 there was probably less likely report of disease, resulting in underreporting of COVID-19 cases and therefore mortality rate should be less estimated. Our methodology lies in its ability to detect epitope features that could raise a red flag in the emergence of new variants from distinct spike protein clusters able to potentially evade immune responses induced by vaccination or previous infections. Nonetheless, the primary conclusion of the *in silico* analysis requires, indeed, extensive wet laboratory experiments. Also, the identifying B and T cell-specific epitopes is crucial for immunotherapeutic and prophylatics approaches.

## Acknowledgments

We would like to thank the Núcleo de Processamento de Alto Desempenho da Universidade Federal do Rio Grande do Norte-NPAD/UFRN for allowing us to access their computer facilities. We also gratefully acknowledge the following authors from the originating laboratories listed on Supplemental Material 1 for obtaining the specimens as well as the submitting laboratories where the SARS-CoV-2 spike glycoprotein sequence data were generated and shared via GISAID on which was the bases for this Supplemental Material 1.

The work was supported in part by a grant from Instituto Todos pela Saúde (ITpS) to S.M.B.J. The funders had no role in the design of the work nor the interpretation of the data.

J.F.R.N., D.G.T., D.C.S.C and S.M.B.J. Conceptualization, Data curation, Formal analysis, Investigation, Project administration, Supervision, Writing – original draft, Writing – review & editing. J.F.R.N. and D.G.T., Methodology, Validation. S.M.B.J., Funding acquisition.

The authors report no conflict of interest.

